# OASIS evaluation of the French surveillance network for antimicrobial resistance in diseased animals (RESAPATH): success factors at the basis of a well-performing volunteer system

**DOI:** 10.1101/2021.04.07.438805

**Authors:** R. Mader, N. Jarrige, M. Haenni, C. Bourély, J.-Y. Madec, J.-P. Amat, on behalf of EU-JAMRAI

## Abstract

Antimicrobial resistance is a One Health issue requiring the development of surveillance systems in the human, environmental and animal sectors. In Europe, the surveillance of antimicrobial resistance on zoonotic pathogens and indicator bacteria in healthy food-producing animals has been implemented on a legal basis, while countries are also expected to extend their surveillance to diseased animals in the frame of national action plans. In this context, evaluating existing antimicrobial resistance surveillance systems in veterinary medicine is important to improve systems in place, but also to help other countries learn from these experiences, understand success factors and anticipate challenges. With this aim, the French surveillance network for antimicrobial resistance in bacteria from diseased animals (RESAPATH) was evaluated using the OASIS assessment tool. Key performance factors included (i) a strong and inclusive central institutional organization defining clear and well-accepted surveillance objectives, scope and procedures, (ii) strong skills in epidemiology and microbiology and (iii) a win-win approach enabling the volunteer participation of 71 field laboratories and where a free annual proficiency testing plays a pivotal role. The main area of improvement of RESAPATH was its time-consuming data management system.

## Introduction

Antimicrobial resistance (AMR) is a threat to our societies, accounting for an estimated 700000 deaths per year [1]. To tackle AMR, a global action plan was endorsed at the Sixty-eighth World Health Assembly, which underscores the need for an effective “One Health” approach involving coordination among numerous international sectors and actors, including human and veterinary medicine [2]. In line with this plan, countries are urged to develop One Health AMR surveillance systems. This was further emphasised in the European Union (EU) One Health Action Plan which states that “*a comprehensive, collaborative and coordinated collection and analysis of data from multiple domains, i*.*e. a One Health AMR surveillance system, is therefore essential*” [3].

In Europe, a coordinated and programmed AMR surveillance system has been implemented in the animal sector with the perspective to protect consumers from AMR transmission through the food chain. This EU harmonized AMR monitoring refers to zoonotic pathogens and indicator bacteria recovered from meat samples at retail and healthy animals at slaughterhouses. It does not, however, cover AMR monitoring in veterinary clinics and farms where antimicrobials are effectively used. Therefore, similarly to most - if not all - systems in human medicine, monitoring AMR in diseased animals is of utmost importance in the context of a national action plan. The multiple benefits of AMR monitoring in animal bacterial pathogens are, according to the World Organisation for Animal health (OIE), to:

– *“detect emerging resistance that may pose a concern for animal and human health;*
– *detect changes in susceptibility patterns;*
– *provide information for risk analysis;*
– *provide data for veterinarians to inform their treatment decisions;*
– *provide information for epidemiological studies and trend analysis”* [4].

In this context, the EU Joint Action on Antimicrobial Resistance and Healthcare Associated Infections (EU-JAMRAI) aims to assess existing surveillance systems of AMR in diseased animals in the EU, as part of a larger objective to study the feasibility of a coordinated surveillance of AMR in diseased animals in Europe. Here, we report the strengths and limitations of the French surveillance network for AMR in bacteria from diseased animals (RESAPATH), which is the oldest surveillance system of its kind in Europe (set up in 1982) at least regarding livestock [5]. This work aims to provide valuable information for the implementation of similar systems in other countries and to highlight the multiple benefits of evaluating AMR surveillance systems.

## Material and Methods

### The RESAPATH network

RESAPATH has a steering committee composed of representatives of public and private diagnostic laboratories, the ministry in charge of Agriculture, veterinary professional organizations, as well as microbiologists and epidemiologists from the French Agency for Food, Environmental and Occupational Health & Safety (ANSES). It is coordinated by two laboratories of ANSES, located in Ploufragan-Plouzané-Niort and Lyon, France.

RESAPATH performs “passive” (or “event-based”) phenotypical AMR surveillance in cattle, sheep, goats, swine, chickens, turkeys, rabbits, fish, horses, dogs, cats and exotic animals. In 2017, it was composed of 71 volunteer public or private veterinary diagnostic laboratories collecting resistance data for 56 286 isolates. From 2008 to 2017, it experienced an important development, with an increase of 39% in the participating laboratories and 211% in the data collected [6]. RESAPATH became an important component of the French National Action Plan (NAP) to tackle AMR in the animal sector, so-called ECOANTIBIO^2^ (2017-2021) [7]. The objectives of RESAPATH, as defined in a mutual agreement between participating laboratories and ANSES, are to:

– follow AMR trends in pathogenic bacteria of animals;
– collect and store a panel of isolates for in-depth molecular investigations;
– provide solid technical and scientific support to field laboratories;
– enable comparisons of animal / human AMR data through the French national observatory for epidemiology of bacterial resistance to antimicrobials (ONERBA), to which RESAPATH is federated.

The mutual agreement also defines the roles and duties of the coordination team (ANSES) and participating laboratories.

On the one hand, all member laboratories are required to perform antimicrobial susceptibility testing (AST) by disk diffusion according to the French norm NF U 47-107 [8] and to interpret AST results according to the veterinary guidelines of the Antibiogram Committee of the French Society of Microbiology (CA-SFM) [9]. They are also required to send to ANSES all AST data (antibiotics tested and inhibition diameters) together with appropriate anonymized metadata (geographical origin, animal species, age category, specimen, disease and bacterial species) every three months using a dedicated Excel® template. Upon request, laboratories also provide bacterial isolates with specific phenotypical resistance profiles for further molecular investigations (ANSES bearing shipping costs).

On the other hand, ANSES is committed to organize and financially support an annual proficiency testing (PT) and continuous scientific and technical support to laboratories on the AST technique and results’ interpretation. ANSES is also responsible for the analysis of surveillance data and in charge of communication activities such as (i) editing a publicly-available annual surveillance report, (ii) managing the RESAPATH website (https://resapath.anses.fr/), (iii) issuing a regular newsletter to the network and (iv) organizing an annual one-day RESAPATH meeting where all participating laboratories and other partners are invited.

### Evaluation method

The OASIS tool is a generic method to evaluate any kind of surveillance system and was used to evaluate RESAPATH [10]. This qualitative method was developed in 2010-2011 to perform standardized, detailed and comprehensive evaluations of the organization and operations of surveillance systems. It is the reference method used by the French platform for epidemiological surveillance in animal health [11]. OASIS is based on an evaluation grid composed of 78 criteria which are evaluated from 0 to 3 and accompanied by commentaries which include justifications for attributed scores and recommendations of improvement. It is commonly acknowledged that comments are of great importance and complementary to scores to finely assess the performance of a surveillance system. OASIS outputs are displayed in the form of three complementary figures on (i) ten functional sections (defined according to the structure and activities of a surveillance system), seven critical control points (CCPs) which are the operations at which control can be applied and ten attributes, i.e. measurable characteristics such as representativeness or timeliness which indicate the system’s quality. In these figures, results are indicated as proportions of the maximum possible score.

Once the RESAPATH steering committee approved the principle of an OASIS evaluation, a joint team was built, composed of two external assessors (not involved in RESAPATH activities) and two internal assessors (members of the RESAPATH coordination team) (Table 1). This team conducted a series of semi-open interviews from 11 June 2018 to 10 July 2018, either face to face or by phone, with a panel of 23 representative partners of RESAPATH (Table 2). Most interviews lasted between 1 hour and 1.5 hours. Following the interviews, the evaluation team pre-filled the OASIS grid (including commentaries). Then, the grid was reviewed by a group composed of seven representative partners of RESAPATH (Table 3) and the four assessors during a full-day meeting on 17 July 2018.

**Table 1:** External and internal assessors of the OASIS evaluation of RESAPATH in 2018.

| Name | Position (at the time of evaluation) |
| --- | --- |
| Rodolphe Mader<br>(external assessor) | Epidemiologist at the Antimicrobial Resistance and Bacterial Virulence Unit, ANSES, Laboratory of Lyon. Not involved in RESAPATH activities. |
| Jean-Philippe Amat<br>(external assessor) | Epidemiologist at the Epidemiology and Support to Surveillance Unit, ANSES, Laboratory of Lyon, not involved in RESAPATH activities. |
| Marisa Haenni<br>(internal assessor) | Microbiologist, deputy head of the Antimicrobial Resistance and Bacterial Virulence Unit, ANSES, Laboratory of Lyon, member of the RESAPATH coordination team and member of the scientific board of the French National Observatory for Epidemiology of Bacterial Resistance to Antimicrobials (ONERBA). |
| Nathalie Jarrige<br>(internal assessor) | Epidemiologist at the Epidemiology and Support to Surveillance Unit, ANSES, Laboratory of Lyon, member of the RESAPATH coordination team. |

ANSES: French Agency for Food, Environmental and Occupational Health & Safety.

**Table 2:**
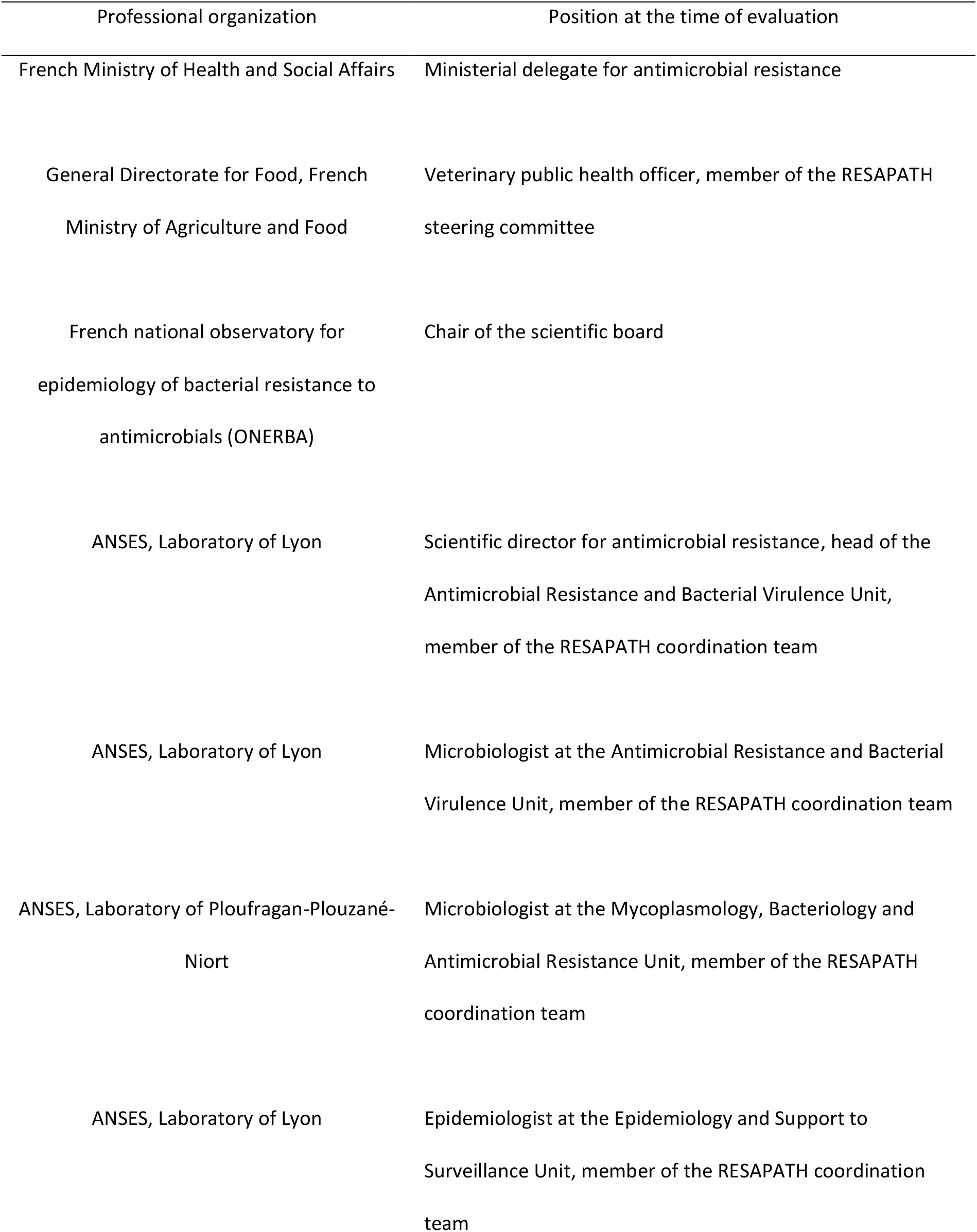
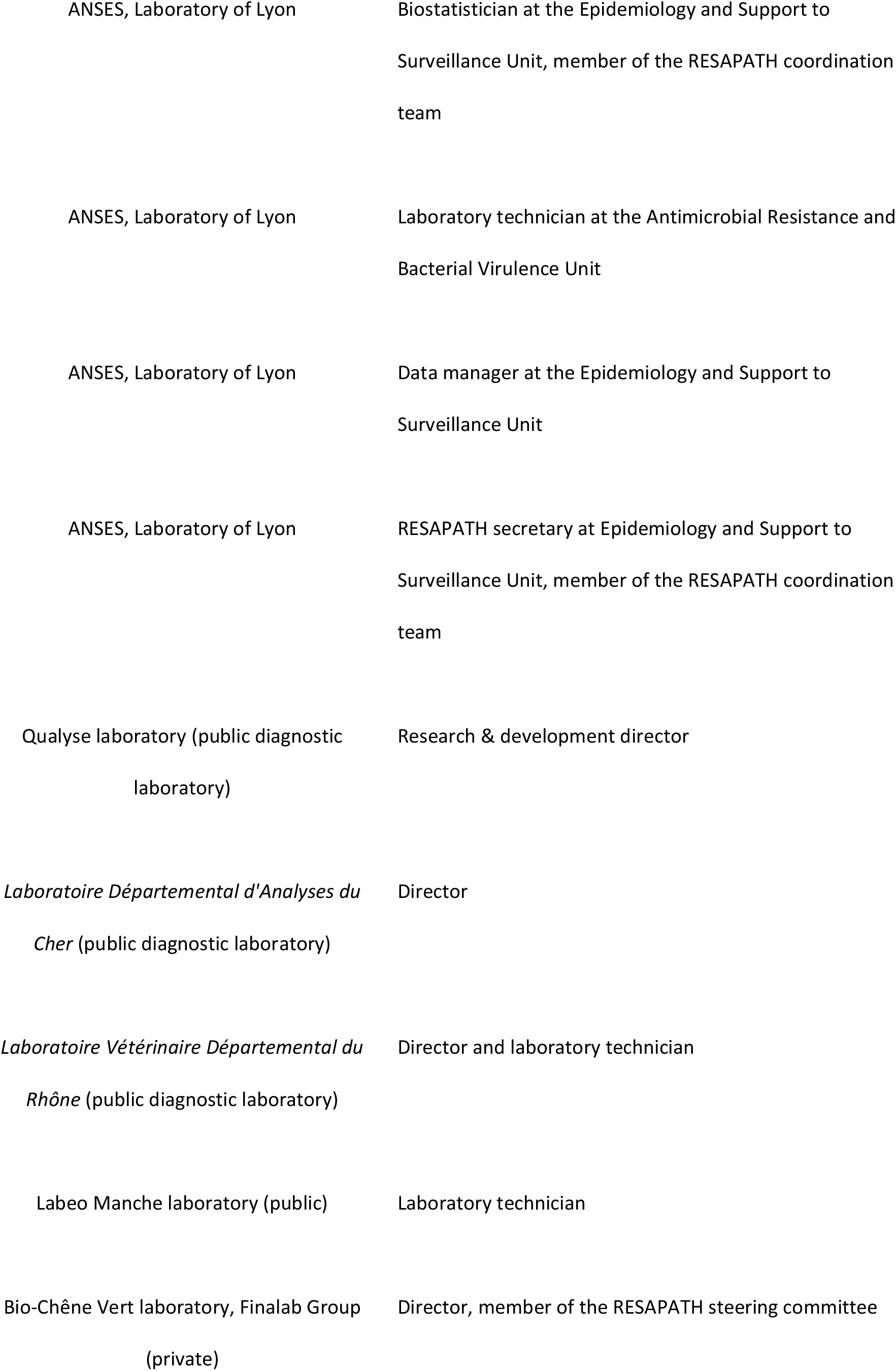
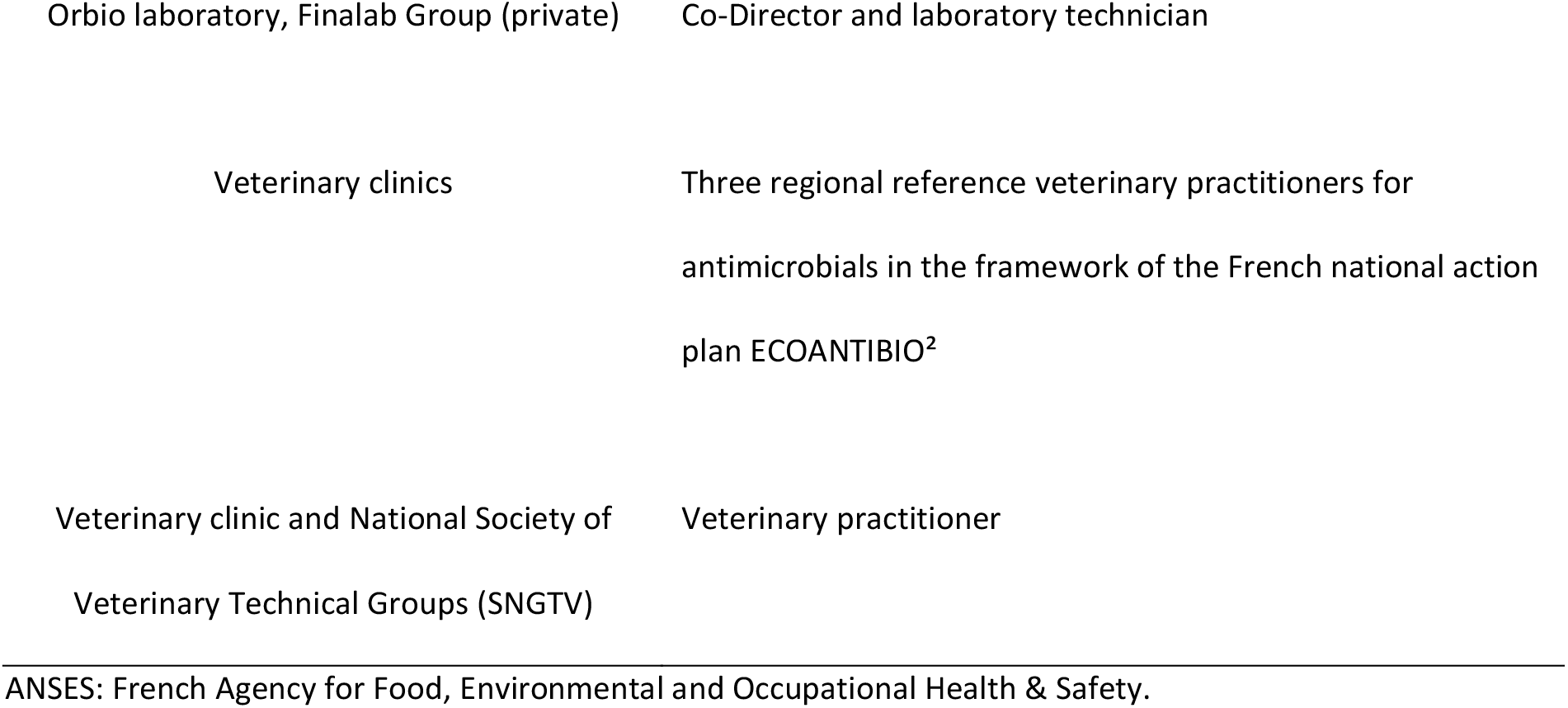
Professional organizations and positions of the 23 RESAPATH partners interviewed during the OASIS evaluation of RESAPATH in 2018.

**Table 3:** Professional organizations and positions of the seven members of the evaluation grid review group (in addition to the four assessors) during the OASIS evaluation of RESAPATH in 2018.

| Professional organization | Position at the time of evaluation |
| --- | --- |
| General Directorate for Food, French<br>Ministry of Agriculture and Food | Veterinary public health officer, member of the RESAPATH<br>steering committee |
| ANSES, Laboratory of Lyon | Scientific director for antimicrobial resistance, head of the<br>Antimicrobial Resistance and Bacterial Virulence Unit,<br>member of the RESAPATH coordination team |
| ANSES, Laboratory of Lyon | Epidemiologist at the Epidemiology and Support to<br>Surveillance Unit, member of the RESAPATH coordination<br>team |
| ANSES, Laboratory of Ploufragan-Plouzané-Niort | Microbiologist at the Mycoplasmaology, Bacteriology and<br>Antimicrobial Resistance Unit, member of the RESAPATH<br>coordination team |
| ANSES, Laboratory of Ploufragan-Plouzané-Niort | Head of the Mycoplasmaology, Bacteriology and Antimicrobial<br>Resistance Unit |
| ANSES, Laboratory of Lyon | Veterinary public health officer |
| *Laboratoire Départemental Vétérinaire de*<br>*l'Hérault* (public diagnostic laboratory) and<br>ADILVA | Laboratory director, ADILVA representative |
| ANSES: French Agency for Food, Environmental and Occupational Health & Safety. ADILVA: French association<br>of public veterinary diagnostic laboratory managers. |  |

## Results

The three OASIS outputs are presented in Figures 1, 2 and 3 and show high scores, with none of them below 50%. The lowest scores were obtained for the field institutional organization, surveillance tools and communication among functional sections (Fig. 1), sampling and information distribution for CCPs (Fig. 2) and representativeness and timeliness regarding attributes (Fig. 3).

**Fig. 1:**
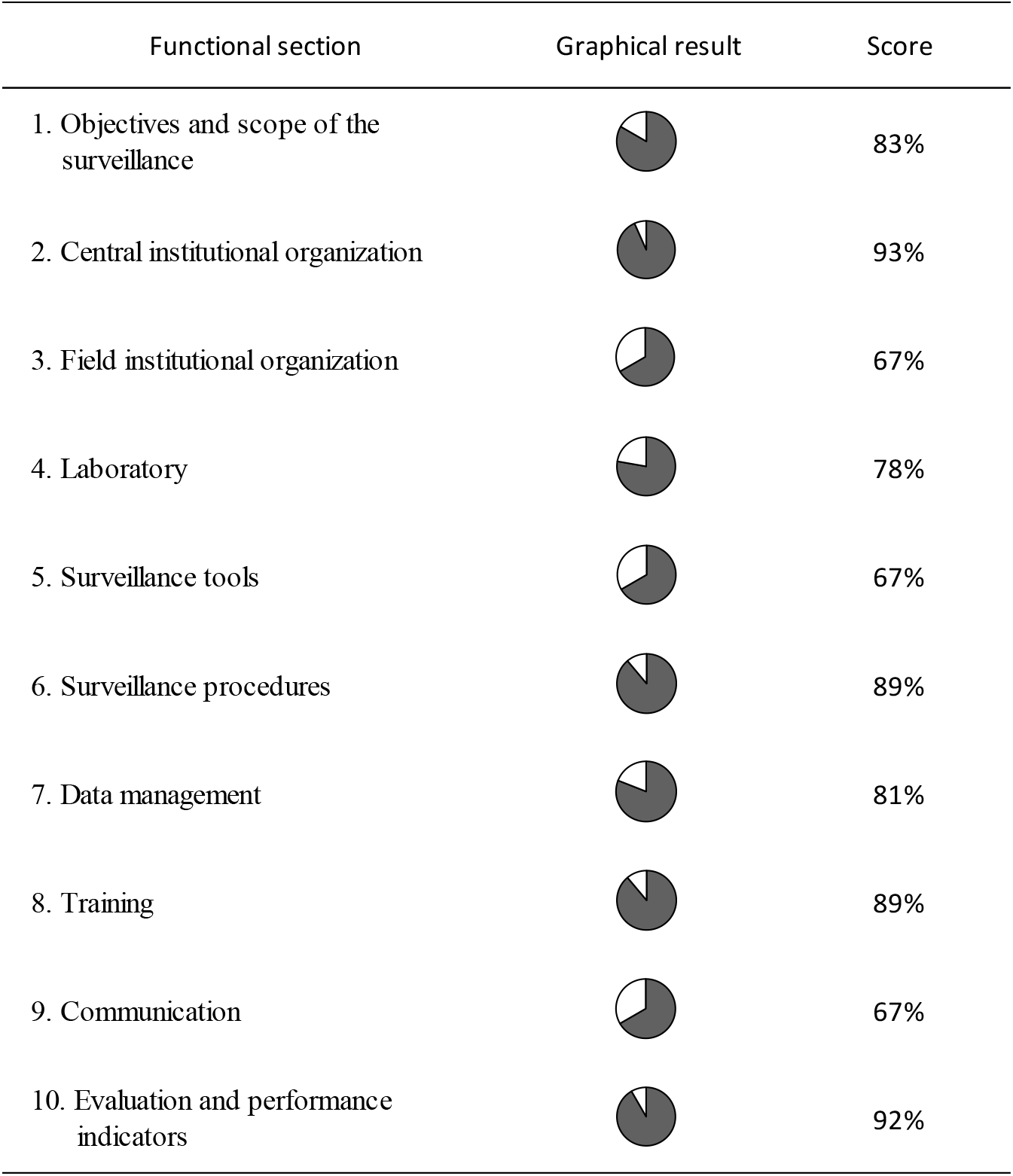
Scores of RESAPATH obtained to the ten OASIS functional sections (indicated in proportions of the maximum possible score) in 2018.

**Fig. 2:**
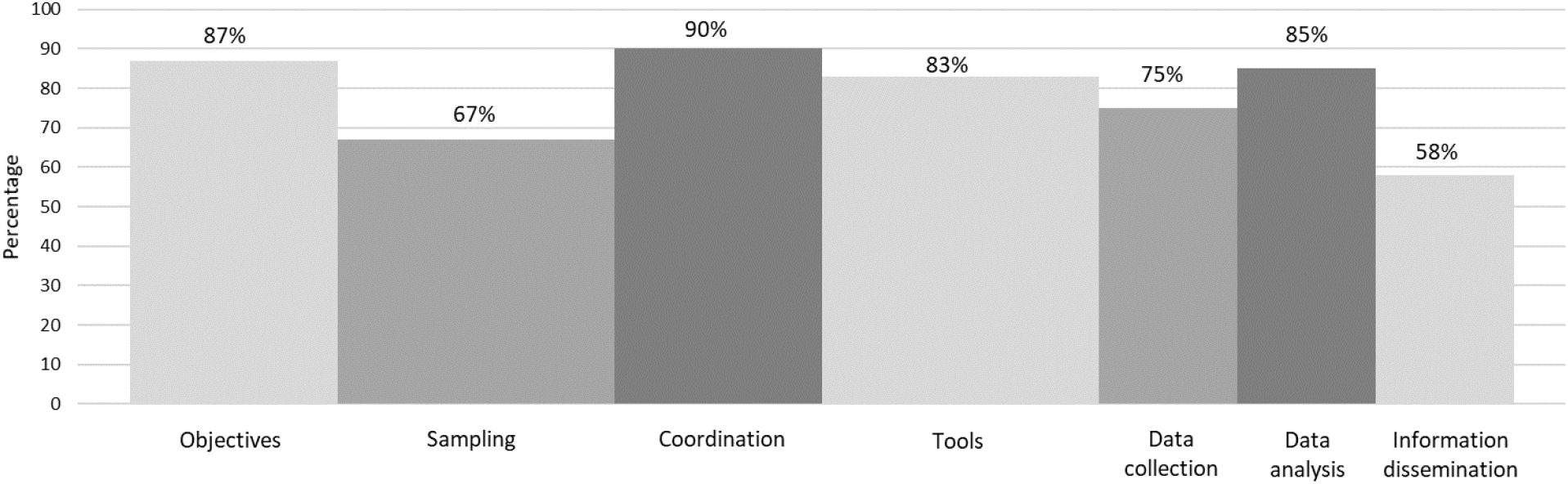
Scores of RESAPATH obtained to the seven OASIS critical control points (indicated in proportions of the maximum possible score) in 2018.

**Fig. 3:**
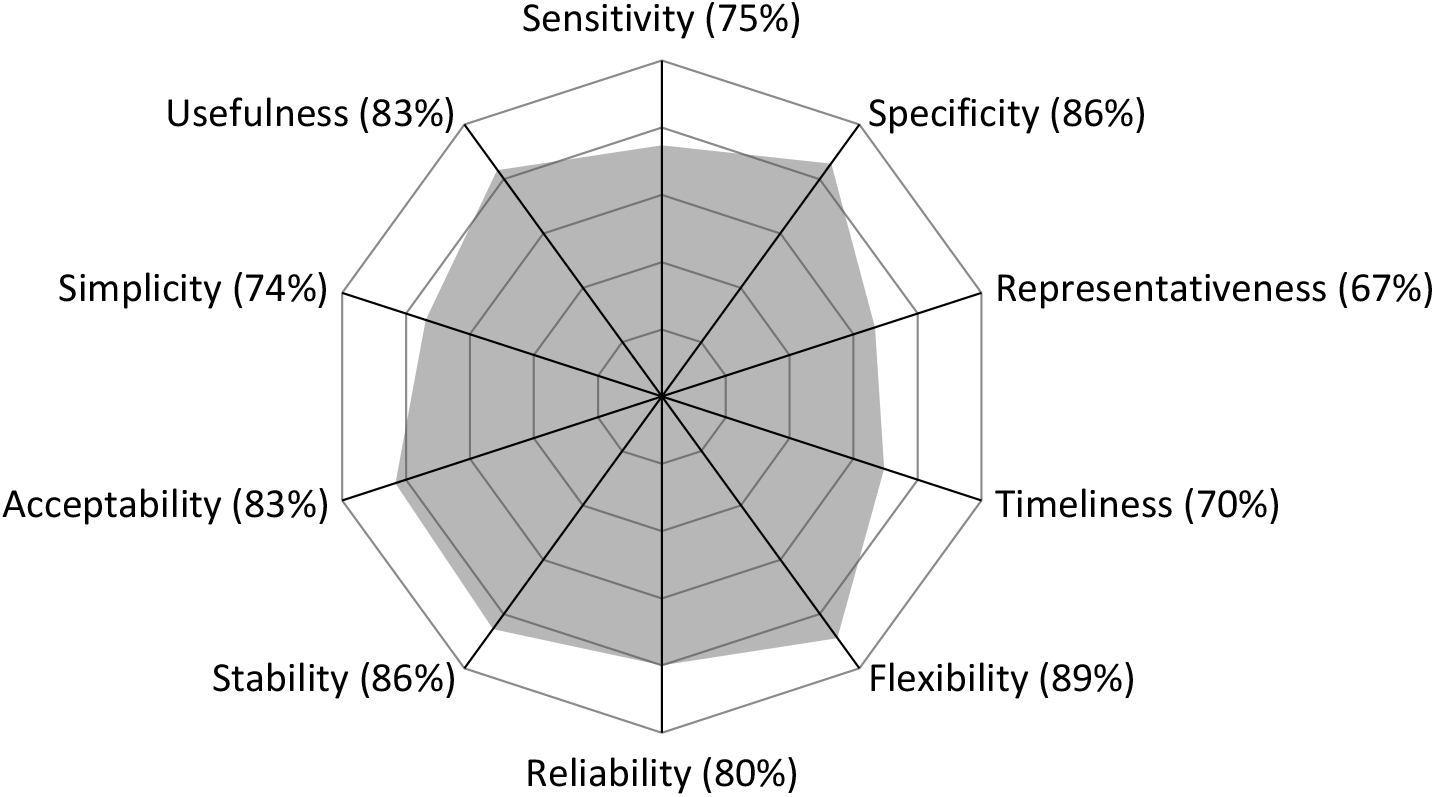
Scores of RESAPATH obtained to the ten OASIS attributes (indicated in proportions of the maximum possible score) in 2018.

The surveillance objectives were considered relevant (Figs. 1 and 2) although some of them were not strictly speaking surveillance objectives and should be reformulated (Table 4). The surveillance scope of RESAPATH is very large since laboratories send all AMR data originating from clinical animal samples. However, limitations in human resources prevent further integration of additional laboratories and have a significant impact on communication and laboratory training activities. It also reduces the proportion of samples subjected to molecular analyses. To enable these developments, an increase in financial and human resources of the coordination team was recommended (Table 4).

**Table 4:**
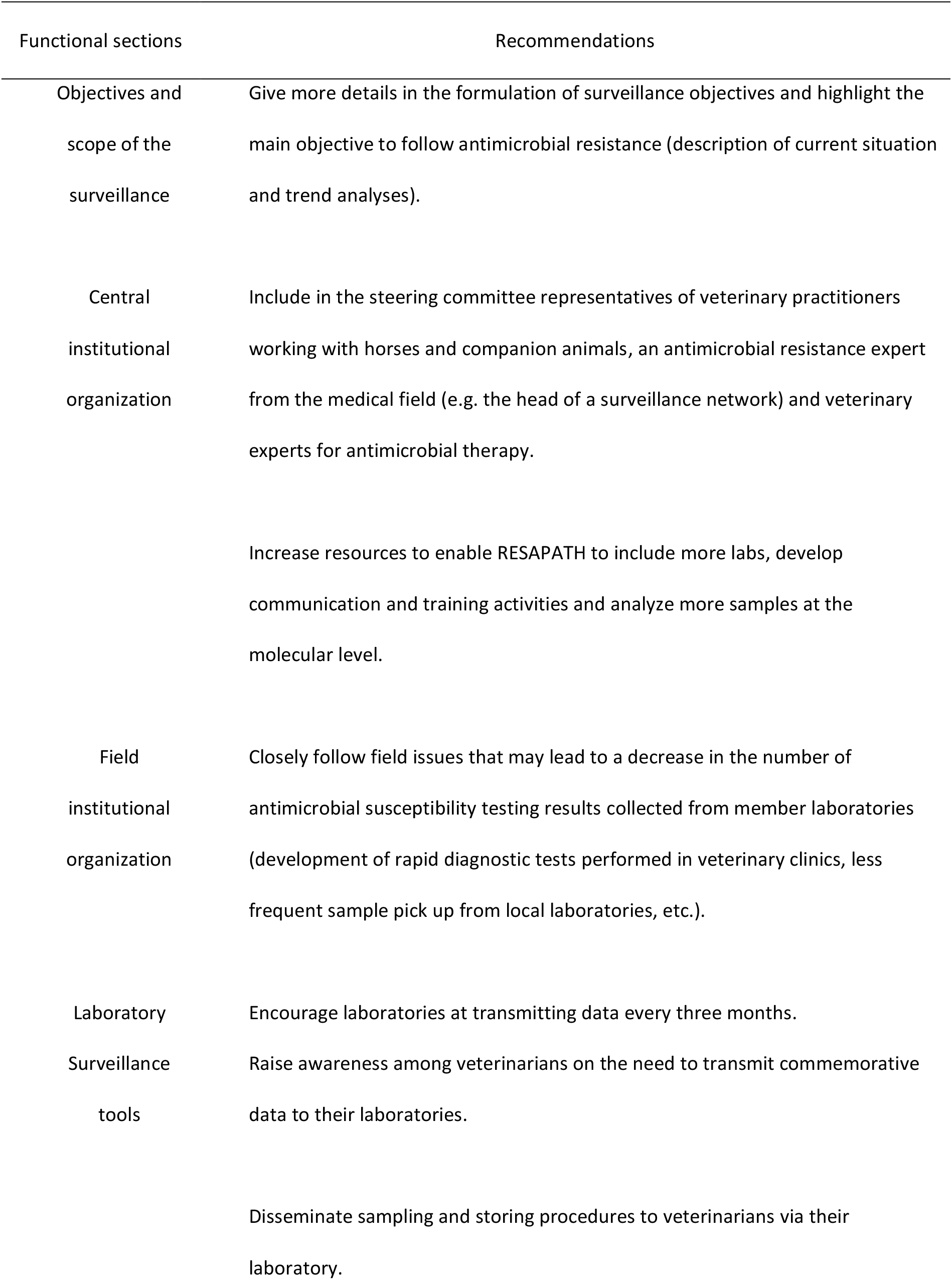
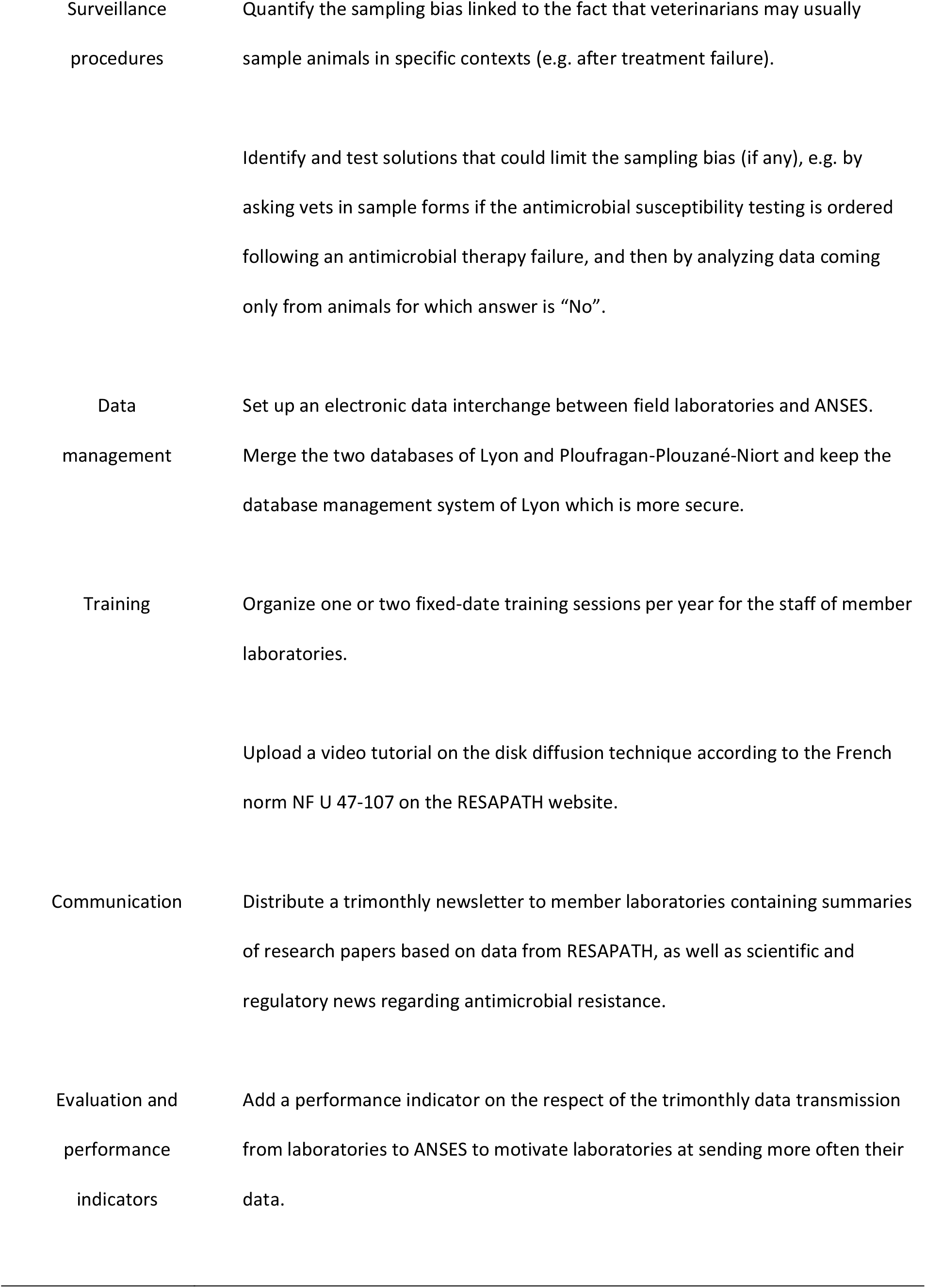
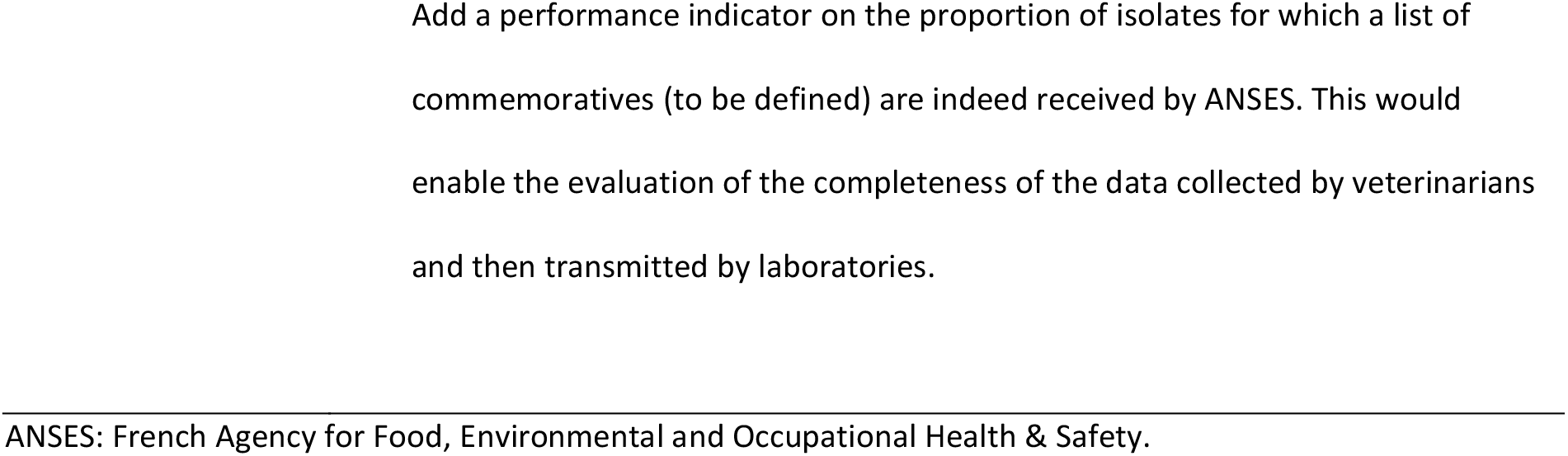
Main recommendations of improvement following the OASIS evaluation of RESAPATH in 2018.

The field institutional organization is composed of veterinary clinicians and laboratories. Thanks to the presence of member laboratories in all French administrative regions, as well as a good geographical overlapping of distributions of animal populations and AST data [12], the geographical coverage and representativeness (Fig. 3) of RESAPATH was considered satisfactory, although not perfect. The sub-optimal score to the field institutional organization (Fig. 1) was also due to financial and practical limitations for ordering an AST (such as delays to get results or less frequent sample pick up from veterinary clinics) and the simultaneous development of rapid AMR diagnostic tests directly performed by veterinarians and therefore not collected by the network. It was recommended to follow these evolutions carefully (Table 4).

The central institutional organization defined simple and adapted surveillance procedures (Fig. 1) participating in the good scores obtained for simplicity and acceptability (Fig. 3). However, the “passive” data collection entails possible sampling biases (Fig. 2) leading to sub-optimal representativeness of pathogenic bacterial populations of animals (Fig. 3). Indeed, animals may be more prone to be sampled in specific circumstances, such as previous treatment failures or chronic infections. Nonetheless, this hypothesis proved invalid in poultry productions where Bourély *et al*. showed that AST were performed in merely all suspicions of infection [13]. Therefore, we recommended to quantify such possible biases outside the poultry sector and suggested a solution to address sampling bias issues (Table 4).

The score for surveillance tools was considered satisfactory (Figs. 1 and 2) but negatively impacted by the lack of inclusion of roles and duties for veterinarians in the mutual agreement. Thus, RESAPATH has no control on the collection, storage and transfer of clinical samples and related commemoratives from veterinarians to laboratories, bringing possible limitations (although never assessed) in terms of data quality, completeness and harmonization. As such, laboratories could disseminate sampling and storing procedures to veterinarians and raise their awareness on the need to transmit commemoratives (Table 4).

Participating laboratories are at the basis of the system. They have strong methodological skills and the disk diffusion method is fully harmonized within the network and checked annually during the annual PT. All laboratories have the possibility to follow a free technical training to improve their skills on the AST method upon demand. These characteristics contributed to high scores at the training and laboratory functional sections (Fig. 1). However, interviews revealed that laboratories can be hesitant at asking for such trainings, so we recommended to organize formal training sessions at fixed dates in parallel to convenient training on sites and to upload a video tutorial on the disk diffusion technique on the RESAPATH website (Table 4).

RESAPATH received a good score on its data management functional section (Fig. 1), but at the price of significant time-consuming processes for both the laboratories and the coordination team. Indeed, many laboratories do not have optimal Information Technology (IT) material. Some have to report manually their AMR data in the RESAPATH Excel® template, while others can make a data extraction from their laboratory information management system, but not in the required format and codification. In the end, the coordination team receives a great diversity of file formats and had to develop a series of semi-automated solutions to convert laboratory files into the appropriate format in order to reduce the workload of data cleaning and harmonization. In addition, most laboratories do not send their data every three months, leading to heavier workloads at some periods. This situation led to a sub-optimal score to data collection (Fig. 2) and timeliness (Fig. 3). However, all laboratories send their data at least once a year, enabling the analysis of the complete RESAPATH database each year. Another weakness is the existence of two RESAPATH databases in two different ANSES laboratories: one in Ploufragan-Plouzané-Niort with data from swine, chickens, turkeys, rabbits and fish and another one in Lyon with data from other animal species, mainly ruminants, equids and companion animals. This leads to a lack of efficiency in processes such as data cleaning, integration or analyses, which are carried out separately. Moreover, the database in Ploufragan-Plouzané-Niort is not as secured as the one in Lyon. Thus, data management was assessed as the main area of improvement for RESAPATH and the main cause for not accepting additional laboratories in the network. The project to set up an electronic data interchange between laboratories and ANSES may solve some of these issues and so has become a recommendation from this evaluation too (Table 4). Specific attention should nevertheless be paid not to exclude small laboratories with limited IT capacities from the network. In the meantime, a more frequent data transmission from laboratories should be encouraged (Table 4). Finally, it was recommended to merge both databases and keep the database management system of Lyon (Table 4).

Despite these weaknesses, RESAPATH manages to perform well in terms of data analyses (Fig. 2), with also good scores to sensitivity, specificity and reliability (Fig. 3). Its capacity to produce multi-disciplinary analyses thanks to skills in bacteriology, epidemiology, biostatistics and IT within the coordination team was considered as a strong asset.

A particularity of RESAPATH, which drives its whole performance, is its capacity to develop a volunteer network of laboratories thanks to a win-win approach. Indeed, RESAPATH is not regulated by law, not supported by dedicated funds and relies on the volunteer participation of all members. According to the interviews, the main motivation for integrating RESAPATH was the so-called “free” and high quality technical support that laboratories receive. This support includes an annual PT which was recognized as a key benefit for laboratories. This PT has the double advantage of enabling laboratories to check and continuously improve their own performance and for ANSES to check the quality of the AMR data collected. Some laboratories also use their participation in this PT as a label of quality for marketing purposes. In turn, laboratories accept to send AMR data and isolates of particular interest, allowing long-term AMR surveillance at the national level and providing material to researchers in microbiology and epidemiology at ANSES. This win-win approach, where ANSES and participating laboratories always need to meet each other’s expectations, is at the heart of the dynamic of RESAPATH and the basis of its strong acceptability, flexibility and stability scores (Fig. 3). Beyond acceptability, the RESAPATH community seems to have even developed a sense of belonging according to our interviews. Moreover, RESAPATH enables its member laboratories to gain a better recognition as indisputable actors in the fight against AMR in the animal sector.

Scores for communication and information distribution were not as high as others (Figs. 1 and 2) due to the absence of any newsletter distributed to the network. Issuing a newsletter on a regular basis is mentioned in the mutual agreement and is expected by laboratories, so we recommended to put efforts on this point (Table 4). However, the assessors also acknowledged the numerous and successful efforts of the coordination team to develop internal and external communication through the organization of an annual RESAPATH meeting, frequent and increasing email exchanges, numerous participations in veterinary, epidemiology and microbiology events, participation in the ONERBA, the public release of annual reports (available in French and English on the RESAPATH website) and scientific articles, as well as an ongoing project to develop a web application (using the Shiny package [14] of R software [15]) to make RESAPATH results more readily accessible to all, especially veterinary practitioners. In the end, we considered that RESAPATH succeeded in maintaining the momentum and motivation of all partners.

Regarding evaluation, RESAPATH received a high score (Fig. 1) thanks to the annual calculation and analysis of relevant internal performance indicators and the undertaking of two OASIS evaluations in 2010 and 2018, despite the fact that limited human resources can prevent the implementation of certain recommendations. It was also suggested to add two performance indicators, one on the respect by laboratories of the trimonthly data sending and another one on the completeness of commemorative data (Table 4).

Finally, RESAPATH proved to be a useful system (Fig. 3), which in turn reinforces its sustainability. An illustration of this is its recent recognition as a key partner of the French NAP to tackle AMR. Indeed, RESAPATH enables the ministry in charge of Agriculture to monitor the efficiency of the NAP and to address some urgent issues, such as in 2015 when a rapid assessment of the spread among animals of the newly discovered plasmid-mediated *mcr-1* colistin resistance gene was required [16]. Veterinarians also benefit from the regular production of epidemiological results on resistance levels that can guide their treatment decisions. Finally, the regular improvement of the performance of diagnostic laboratories leads to the delivery of higher quality AST results to veterinary practitioners with likely clinical and economic impacts.

## Discussion

To our best knowledge, it is the first time that a complete evaluation of an AMR surveillance network in veterinary medicine is published. As such, this study provides valuable information to countries aiming to set up a national AMR surveillance system in diseased animals or to those wishing to improve their current system.

From this evaluation, key success factors for RESAPATH were identified. They included:

– A strong central institution with the inclusion of many relevant actors in the animal sector defining clear surveillance objectives, scope and procedures that meet their expectations;
– Strong skills in epidemiology and microbiology at the central level, including the capacity to complement phenotypical surveillance with molecular surveillance;
– A collaborative win-win approach between member laboratories and the coordination team, leading to the volunteer participation of numerous field laboratories throughout the whole country;
– The provision of free technical support to laboratories including an annual PT enabling the production of high-quality and harmonized AST data;
– A strong internal and external communication with all possible end-users of surveillance data, including in the human sector;
– A continuous self-improvement dynamic through OASIS evaluations, use of performance indicators and organization of laboratory trainings.

However, this evaluation pointed out several areas of improvement. The most important one referred to data management and a lesson to learn would be to always anticipate the possible scalability, a parameter which is often overlooked at the setup of a system. Taking into account the limited IT capacities of laboratories is key to maintaining a strong volunteer network, but this requires a lot of flexibility from the coordination team and can be very time-consuming. Another weakness of RESAPATH relies on possible sampling biases, as a passive laboratory-based surveillance network which does not integrate the sampling stage in its procedures. However, these possible biases were not considered as having a major impact on representativeness in this evaluation. On the other hand, the current organization of RESAPATH brings a lot of simplicity, which is key to its sustainability.

Veterinary medicine is most likely structured similarly in European countries, which makes us believe that passive and volunteer AMR surveillance systems in diseased animals may be built in other places. However, duplication of existing systems should not be the way to go. On the contrary, fine and country-specific analyses and a wide inclusion of all relevant actors are essential to design a well-performing system. RESAPATH shows how participative win-win surveillance networks are of strong value for national authorities. Those systems also trigger highly positive collateral impacts, such as capacity building in veterinary diagnostic through greater skills acquired by all members of the network.

The OASIS tool also proved successful with fruitful exchanges and a strong acceptability of its results and recommendations. At the time of writing, the implementation of some recommendations has already started with the organization of fixed-date training sessions, the ongoing merging of the two databases and the development of the electronic data interchange between laboratories and ANSES.

However, OASIS remains a generic tool and some debates have occurred on the relevance and interpretation of some of its evaluation criteria in the specific case of a passive laboratory-based AMR surveillance system. For example, should the data collection be evaluated from the laboratory stage and/or from the veterinary stage? The review group succeeded in taking consensual decisions, but this can slightly hinder the comparability of subsequent evaluations if such decisions are not consistent in time (this was namely the case for several criteria between this evaluation and the one performed in 2010). The OASIS method has also currently some limitations as it does not enable economic evaluations (e.g. cost benefit analyses) or to investigate multisectoral collaboration, something of particular value for One Health issues such as AMR. To find the most appropriate method, depending on the evaluation question and surveillance attributes to evaluate, assessors may follow the steps suggested in the RISKSUR EVA tool, a framework providing guidance in the planning, implementation and reporting of evaluations [17]. Regarding multisectoral collaboration, a specific tool called ECoSur (https://survtools.org/wiki/surveillance-evaluation/doku.php?id=quality_of_the_collaboration) was recently developed to allow for an in-depth analysis of the organization and functioning of collaboration taking place in a multisectoral surveillance system [18].

Of note, a method specifically dedicated to the assessment of national AMR surveillance systems has been developed since 2015 by the Food and Agriculture Organization of the United Nations (FAO). It was based on the FAO Surveillance Evaluation Tool (SET), itself inspired from OASIS [19], and on the FAO Laboratory Mapping Tool (LMT) [20], adapted to assess the specific issues linked to AMR. This method uses the FAO Assessment Tool for Laboratories and Antimicrobial resistance Surveillance Systems (FAO-ATLASS) [21], which consists in two complementary modules (laboratory and surveillance) covering the key components of a national AMR surveillance system in the food and agriculture sectors. It also includes a Progressive Improvement Pathway (PIP) scoring system, designed to assist policy makers in prioritizing actions for building reliable national AMR surveillance systems for the sectors assessed. As such, conducting an FAO-ATLASS assessment could complement our results by providing a more global picture of the performance of France in terms of AMR surveillance in both animal and environmental sectors.

## Conclusion

Overall, RESAPATH exhibited good scores, proving that a well-performing participative surveillance system of AMR in diseased animals is a realistic option to be included in the frame of a NAP. The thorough description and analysis of its organization and operations led to the identification of key success factors among which (i) a strong and inclusive central institutional organization defining clear and well accepted surveillance objectives, scope and procedures, (ii) strong skills in epidemiology and microbiology and (iii) a win-win approach enabling the volunteer participation of 71 field laboratories and where a free annual PT plays a pivotal role. The OASIS evaluation also allowed to identify areas of improvement and provided a series of recommendations. Some of them have already been implemented by RESAPATH, illustrating the usefulness of such evaluations. In a context where AMR surveillance systems should be set up or improved in veterinary medicine, these results are most likely very helpful to other countries designing or improving their own system. Because of their multiple benefits, such evaluations should be encouraged and their results shared at the European and global levels.

## Acknowledgements

We are very grateful to all the professionals interviewed for this evaluation and/or participating in the full-day review meeting on 17 July 2018. We are also thankful to Pascal Hendrikx for his technical support on the OASIS tool and to Nicolas Keck, Michaël Treilles, Francesca Latronico and Béatrice Mouillé for their inputs regarding the FAO-ATLASS tool.

